# A longitudinal data framework for context-specific genotype-to-phenotype mapping

**DOI:** 10.1101/2025.05.07.652202

**Authors:** Thomas Veith, Richard Beck, Vural Tagal, Tao Li, Saeed Alahmari, Jackson Cole, Daniel Hannaby, John Kyei, Xiaoqing Yu, Konstantin Maksin, Andrew Schultz, HoJoon Lee, Aaron Diaz, Janine Lupo, Issam ElNaqa, Steven Eschrich, Hanlee P. Ji, Noemi Andor

## Abstract

Molecular assays can resolve clonal structure, but they are expensive and typically sparse in time, whereas phenotypic observations such as imaging can be collected frequently but often are not preserved in the context needed for later interpretation. We present [CLONEID](http://dev.cloneid.org/), an event-based framework for organizing clone-resolved phenotypic, molecular, and specimen-context records so that genotype-to-phenotype interpretation can be maintained across time. CLONEID links time-stamped Events, assay-specific Perspectives, and reconciled Identities through structured ingestion, provenance-aware retrieval, and reproducible export, complementing upstream clone-calling methods. In a long-term gastric cancer density-selection experiment, CLONEID linked repeated culture events, growth measurements, and late karyotypic profiling within a shared record, supporting longitudinal interpretation of phenotypic adaptation together with underlying chromosomal state.

---

Tumors and experimental cancer models often contain multiple co-existing clones whose phenotypic behavior and molecular composition can differ or change over time [1, 2, 3, 4, 5, 6, 7, 8, 9, 10]. This intra-system heterogeneity creates a particularly informative setting for genotype-to-phenotype mapping, because related clones can differ in molecular state and behavior while still sharing much of their broader biological and environmental background. Here, we use clone operationally to mean a cell population inferred to share a common recent lineage and sufficiently similar underlying state to be treated as a single evolving unit across measurements, space, and time. Many methods infer clonal structure from sequencing and related assays [11, 12, 13, 14, 15]. By contrast, far fewer frameworks address the longitudinal data-integration problem: storing clone-resolved measurements together with the event and specimen context needed to connect phenotypic observations to later molecular readouts [16, 17]. We developed CLONEID to address this problem as an event-based architecture and data model for capturing, linking, and reconciling longitudinal clone-resolved data across clinical, in vivo, and in vitro settings.

CLONEID is organized around three core representational primitives—Event, Perspective and Identity—and one first-class measurement layer, Phenotype (Table **??**, Figure 1). Events record time-stamped propagation, sampling and intervention history together with specimen relationships and environmental context, thereby anchoring the history of a specimen in time and context. Perspectives are assay-specific molecular views of the specimen, typically partial and often obtained through destructive measurement, while preserving the provenance needed to interpret them. Because no single Perspective fully captures the machinery that gives rise to phenotype, CLONEID links multiple Perspectives to approximate an inferred clone-level Identity that can be followed across assays and over time. Phenotype refers to directly witnessed properties of the intact system. This includes *phenotype state* at a given event and *behavioral phenotype* inferred across repeated events over time. Molecular measurements, even when dynamic or rate-based, are not phenotype because they do not directly witness the intact system’s behavior or state; they provide assay-specific molecular perspectives on it. By linking phenotype directly to key events, CLONEID is designed to reduce a common failure mode in longitudinal studies: collecting molecular profiles without the contemporaneous phenotypic context needed for genotype-to-phenotype interpretation.

**Figure 1:**
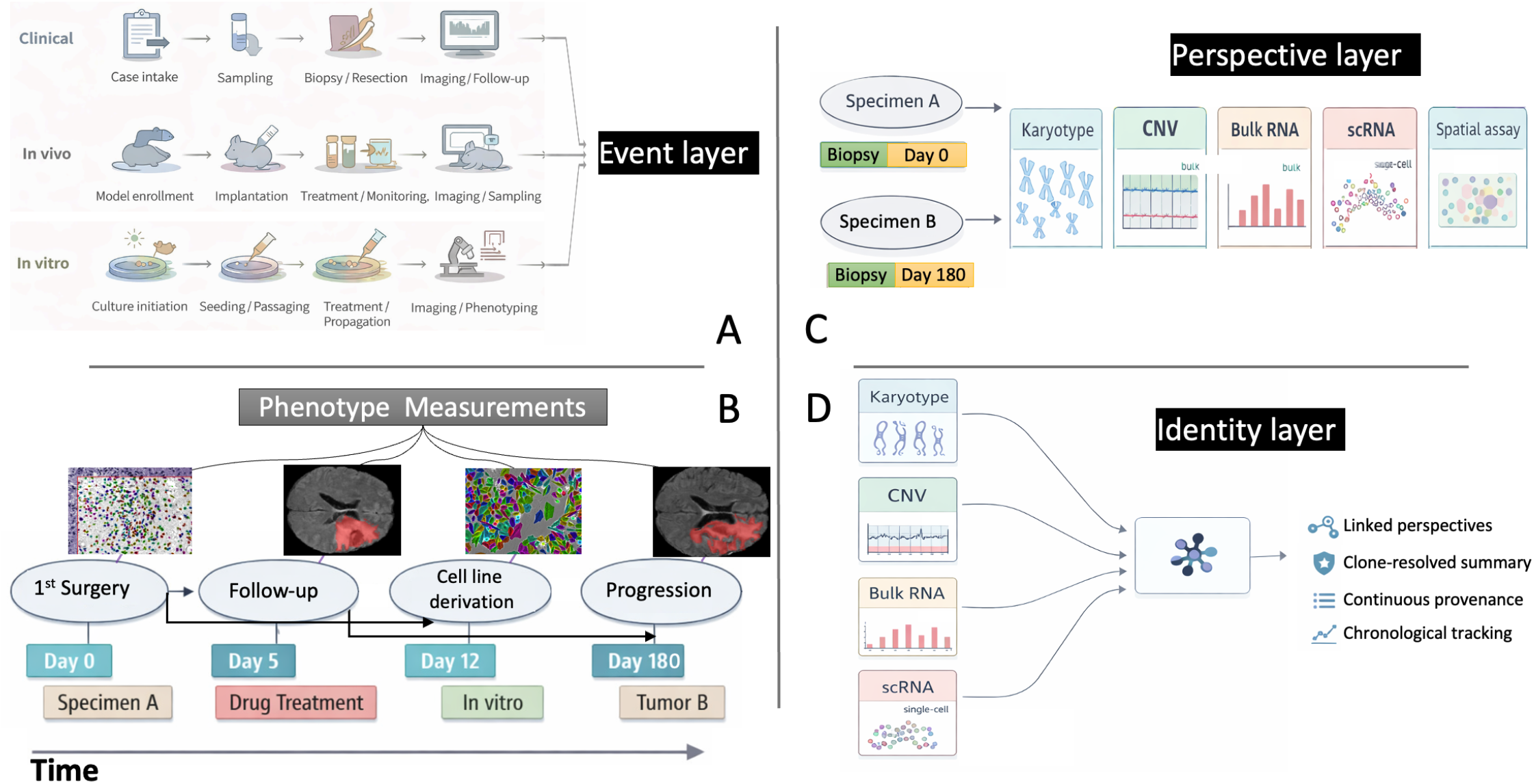
Event-based architecture of CLONEID. (A) The same data model is used across clinical, in vivo and in vitro settings. Time-stamped **Events** capture specimen relationships, interventions and environmental context. (B) Longitudinal phenotype measurements anchor every event. (C) Assay-specific molecular readouts are represented as **Perspectives**, which preserve what was measured together with the upstream data required for provenance. (D) Multiple Perspectives are reconciled into an inferred **Identity** across assays and time. Together, this design enables integrated, queryable and reproducible retrieval of longitudinal genotype–phenotype records.

Here we present the CLONEID architecture and its implementation for ingestion and retrieval of event-linked phenotypic and molecular records, with an emphasis on provenance-aware integration and reproducible access to longitudinal datasets. We highlight the system using a primary end-to-end demonstration in a long-term density-dependent selection experiment in gastric cancer cells, where repeated culture events and growth measurements informed subsequent karyotypic profiling and enabled longitudinal linkage of phenotypic adaptation to underlying chromosomal variations.

An **Event** records “what happened, when, and to which specimen,” spanning propagation, sampling, and interventions across in vitro, in vivo, and clinical timelines (Fig. 1A). Each Event stores a time stamp, event type, specimen relationships, and contextual metadata, thereby anchoring downstream phenotypic and molecular records. CLONEID uses a common transition structure across study settings: genesis Events may have no parent, whereas non-genesis Events are linked to a precursor record so that entry into a new physical or contextual state remains connected to the history from which it arose. This rule ensures that Events record the state associated with the location being entered or exited, rather than merely the chronological order of records. This matters biologically because abrupt context changes, such as transplantation-like transitions, can alter the selective constraints acting on a clone and therefore require especially explicit contextual linkage. In practice, this particularly motivates Event creation when cells relocate into a new physical environment such as seeding, transfer, sampling, biopsy, resection, implantation, or intervention, but Event creation can also be justified by phenotype collection alone, even when no major biological transition has occurred. Because the same Event structure is used across clinical, in vivo and in vitro records, CLONEID represents these settings within a single scenario-neutral architecture (Fig. 1A).

Phenotypes describe the state or behavior of cells or cell populations while the system remains intact and interpretable through its event history. In CLONEID, phenotype includes *phenotype state*, i.e. state at a given Event or timepoint, and *behavioral phenotype*, i.e. a temporal pattern that becomes interpretable only across repeated Events. Behavioral phenotype, by contrast, is inherently temporal and is inferred from repeated Event-linked evidence across time. Event-linked images, cell counts, and related non-molecular records provide observational evidence for phenotype state and also document that the Event occurred. Event-associated observational records, including imaging data, are converted into derived longitudinal measurements that describe phenotype such as segmentation-based features and related image-derived readouts within the same record structure. Phenotypes are by definition holistic, they capture how the intact system as a whole behaves or the state it is in.

In contrast to Phenotypes, Perspectives are assay-specific and inherently partial readouts: each captures only one bounded view of the underlying biological machinery at a particular sampling moment. **Perspectives** (Fig. 1C) can store sample-level summaries, clone-level profiles and individual-cell measurements, while preserving the upstream data needed for provenance and reproducibility. CLONEID can retain both clone-resolved summaries and the underlying measurements from which they were derived, making each Perspective traceable, reproducible and available for downstream feature computation.

Because no single Perspective exhausts what a cell is, multiple Perspectives need to be reconciled to infer **Identity**, the clone-level object that persists across contexts and gives rise to different phenotypes (Fig. 1D). This distinction is central to CLONEID’s larger goal of enabling genotype-to-phenotype mapping. Whereas a `Perspective` records what was measured in a specific assay, an `Identity` records the inferred clone-level object generated by linking one or more Perspectives that are considered consistent with the same underlying clone across assays, samples or time points. This reconciliation is encoded as an explicit relational layer: Perspectives remain assay-specific records, and Identity stores the links that group them into a shared clone representation while preserving their original provenance.

These linked records are stored in an SQL relational database, accessible through two complementary interfaces: a web portal for interactive browsing, upload and export of longitudinal records, and an R package (*thecloneredesignlab/cloneidR*) for programmatic ingestion, Perspective registration, Identity reconciliation, querying and visualization. Both interfaces operate on the same event-based data model, allowing event-linked phenotypic records and assay-specific molecular Perspectives to be ingested, reconciled and retrieved within a common schema. To support portable and reproducible use, CLONEID preserves provenance at the level of both Events and Perspectives and provides context-bundled export of selected record subsets. In the current implementation, exported datasets retain the linkage between event history, phenotypic observations and downstream molecular records, allowing downstream analysis without requiring direct access to the full database.

We used a long-term density-selection experiment in SNU-668 gastric cancer cells as a primary demonstration because it exercises all four pillars of CLONEID in a single setting: repeated Events, longitudinal phenotype capture, late molecular Perspectives and their reconciliation within a shared experimental history. Cells were maintained for more than six months under either low-confluence (r-selection) or high-confluence (K-selection) conditions, generating event-linked growth measurements and a late karyotypic readout for each adaptation history (Fig. 2). Growth trajectories diverged by selection regime (Fig. 2D), and late karyotypic profiles distinguished adaptation histories, including induction and expansion of whole-genome doubled cells in an r-selection regime (Fig. 2E,F). Together, these data suggest that specific karyotypic states are selected under distinct density conditions and contribute to context-specific phenotypic adaptation. Additional appendix analyses show that the same record structure also supports distinct downstream study designs, including prediction of long-term adaptation and cellular plasticity from transcriptomic snapshots (Fig. S4) and mathematical inference of patient-specific karyotype fitness landscapes from multimodal longitudinal glioblastoma records (Fig. S5).

**Figure 2:**
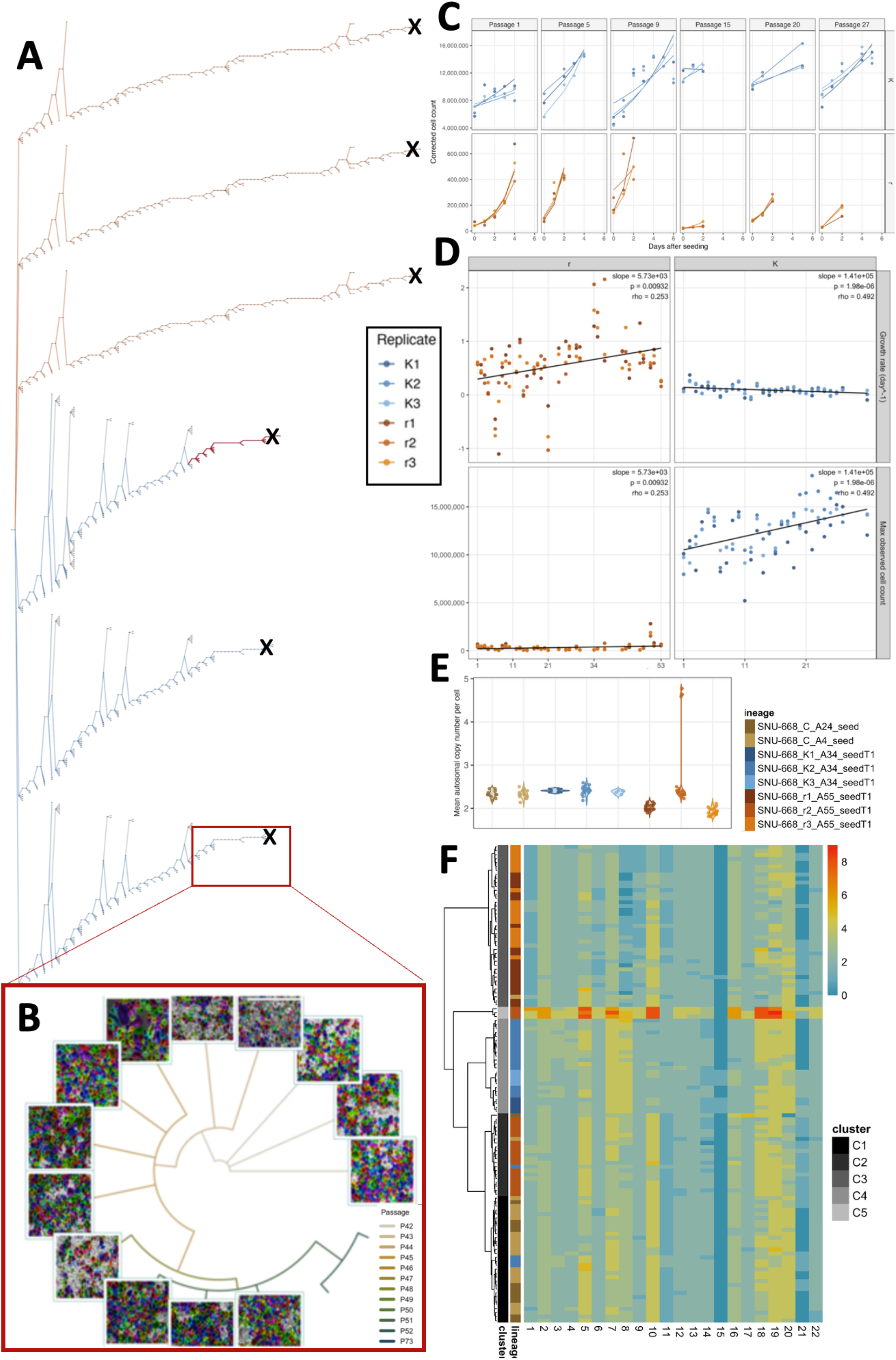
CLONEID links event history, growth dynamics and karyotypic state in a longitudinal density-adaptation experiment. (A) Event history of the full (r/K) adaptation subtree rooted at the parental SNU-668 lineage is shown to place all downstream trajectories in their common ancestry context. Karyotyping timepoints are marked by ‘X’. The branch leading to the image-bearing subset in (B) is highlighted. (B) A circular view of a K-selected sublineage shows representative microscopy overlays together with the corresponding event history and passage progression. (C-D) **Phenotypes**. (C) Daily cell counts and fitted growth curves are shown for matched passages across replicates adapting to sparse (r-selection) or dense (K-selection) population conditions. (D) Growth rate and maximum population size are summarized across passages for all r- and K-selected replicates. (E,F) **Perspective and Identity** profiles distinguish adaptation histories in SNU-668. (E) Mean autosomal copy number per cell is shown as a genome-derived surrogate for ploidy across late control, r-selected, and K-selected specimens. (F) Cell-level karyotypic Perspectives resolve recurrent, condition-associated Identity classes across independent replicates. Code to recreate this figure from data available through dev.cloneid.org is available at https://github.com/thecloneredesignlab/cloneidR/tree/master/analysis.

As of this publication, the CLONEID database contains more than 6,400 event-anchored specimen records organized into 17 rooted lineages across clinical, in vivo and in vitro histories. Data types range from bulk and single-cell DNA/RNA-derived molecular perspectives to image-derived phenotypes. More than 30,000 microscopy, digital pathology and medical images are stored with their associated event context and specimen relationships, along with molecular profiles from over 100,000 individual cells (Fig. S3A). We envision a future in which structured phenotypic data are shared alongside genomic and transcriptomic data as a routine part of cancer research, including manuscript submission and grant-supported data sharing. CLONEID is designed as a step toward that future by providing an event-based framework that links phenotypic measurements to specimen history, assay-specific molecular views and inferred clone identity with preserved provenance. By making these data easier to capture, organize and retrieve across clinical, in vivo and in vitro settings, CLONEID can help turn longitudinal phenotype–genotype records into reusable scientific assets for reproducibility, cross-study integration and future AI-enabled interpretation.

**Table 1:** Ontology-oriented and practical definitions of the core CLONEID concepts. The “ontology definition” column states what sort of thing each concept is in the data model. The “practical / user-facing definition” column states how a user should interpret or instantiate that concept in practice.

| Concept | Ontology definition | Practical / user-facing definition | Key relations |
| --- | --- | --- | --- |
| **Event** | A time-indexed occurrence in the history of a biological system that changes, samples, observes, or otherwise contextualizes the state of a specimen or study entity, and that serves as the anchor for linked evidence, metadata, and relationships. | Create an Event whenever the system enters a new physical or contextual state, and also whenever a directly witnessed property of the intact system is recorded in a way that will matter for reconstructing phenotype over time. An Event answers: <u>what happened, when, and to which specimen?</u> | Anchors event-linked evidence from the intact system and **Perspective** records; non-genesis Events may link to precursor Events; repeated Events support instantiation of **Phenotype state** and derivation of **Behavioral phenotype** across time. |
| **Phenotype** | A first-class layer describing directly witnessed, system-level state or behavior of an intact living system in context. Phenotype functions as the explanandum: it arises from molecular interactions but is not identical to molecular readouts. This includes <u>Phenotype state</u>, i.e. directly witnessed state at a particular Event, and <u>Behavioral phenotype</u>, i.e. directly witnessed pattern inferred across repeated Events over time. | Use **Phenotype state** for directly witnessed properties of the intact system at a given Event, such as morphology, confluence, tumor burden, or other event-local state summaries. Use **Behavioral phenotype** for properties that require repeated Events to infer, such as growth or motility rates. Molecular readouts, even when dynamic or rate-based, are not phenotype. | Grounded in linked **Events** and their associated intact-system evidence; contrasted with **Perspectives**, which provide assay-specific molecular views; used together with **Identity** to support genotype-to-phenotype interpretation and prediction. |
| **Perspective** | A bounded measured view of a specimen, subpopulation, or cell at a particular sampling moment and resolution, representing a molecular layer such as genome, transcriptome, proteome, or karyotype while preserving provenance and assay context. A Perspective captures one partial, snapshot-like aspect of the system. | Use a Perspective for molecular-layer readouts such as genomic, transcriptomic, proteomic, or karyotypic profiles. A Perspective records what molecular layer was profiled, in relation to which Event, and at what resolution, but does not by itself determine phenotype. | Linked to **Events**; one or more Perspectives may be reconciled into an **Identity**; records what was <u>measured</u> about a molecular layer, not what was directly witnessed about intact-system behavior or state. |
| **Identity** | An inferred reconciliation object that links one or more Perspectives judged to be consistent with the same clone-level biological unit across assays, samples, or time, while preserving provenance for that inference. | Use Identity to represent the clone-level object followed across multiple Perspectives. An Identity record should preserve which Perspectives were reconciled, how the linkage was established, and what evidence supports the inference. | Inferred from **Perspectives**; through them remains connected to **Events**; used together with **Phenotype** to support genotype-to-phenotype mapping. |
| **Clone** | An operational biological unit inferred to share recent lineage and sufficient continuity of underlying state to be treated as one evolving unit across measurements, samples, or time. | A clone is not assumed to be observed directly in full. Rather, it is the unit CLONEID attempts to track and approximate through linked Events, Perspectives, Identities, and phenotypes. | Biological target approximated by **Identity**; partial molecular views captured as **Perspectives**; directly witnessed state and behavior summarized through **Phenotype**. |
| **Specimen / study entity** | The material or case-level bearer of state that participates in Events and from which directly witnessed properties and assay-specific records are obtained. Depending on workflow, this may be a patient case, animal model, lineage, sample, or culture-derived entity. | Use this concept for the thing whose history is being organized. In a clinical workflow this may read as a case or specimen; in vivo as an implanted or sampled system; in vitro as a lineage or culture-derived record. | Events occur <u>to</u> or <u>for</u> a specimen / study entity; repeated event-linked intact-system evidence on the specimen supports **Phenotype**; assay-specific molecular measurements are captured as **Perspectives**. |
| **Event-linked evidence** | A provenance-preserving record attached to a specific **Event**, documenting what was observed, measured, or derived at that point, without itself being a phenotype, perspective, or identity. | Use for images, segmentations, counts, annotations, QC outputs, and other event-attached records that support later interpretation. | Anchored to **Event**; may support **Phenotype** or **Perspective**; may contribute evidence used in **Identity** inference. |

## Online Methods

### Software architecture and implementation

CLONEID is implemented as a three-tier system consisting of (i) a user-facing web portal for visualization, upload and export, (ii) a core functionality layer (implemented in Java in the current deployment) for ingestion, linkage and reconciliation of records, and (iii) an SQL database that stores Event-linked phenotypic measurements together with assay-specific molecular Perspectives and their associated meta-data. Uploaded data are managed within an access-controlled warehousing model. Phenotypic image data can be processed using version-controlled analysis applications (for example, Cellpose [18]) so that derived measurements are reproducible and traceable to the originating uploaded observations.

### Demonstration dataset: long-term density-selection experiment

To demonstrate CLONEID in an end-to-end setting, we used a long-term density-selection experiment in the SNU-668 gastric cancer cell line. Cells were cultured in RPMI-1640 supplemented with 10% fetal bovine serum and 1% streptomycin-penicillin at 37°C. Two density-selection regimes were maintained over more than six months: low-confluence (*r*-selection) and high-confluence (*K*-selection). Three replicates were established per regime. Under *r*-selection, replicates were seeded at 135 cells/cm^2^ and transferred upon exceeding 4 × 10^3^ cells/cm^2^, returning the population to 135 cells/cm^2^ at each seeding. Under *K*-selection, replicates were seeded at 10^5^ cells/cm^2^ and transferred upon exceeding 2 × 10^5^ cells/cm^2^. In CLONEID, each seeding and transfer was recorded as an Event linked to its source specimen and contextual metadata.

### Perspectives and Phenotypes

CLONEID distinguishes Phenotype from Perspective by the type of information represented rather than by the measurement modality used to obtain it. Directly witnessed event-level state summaries belong to the Phenotype layer, whereas event-linked images of intact specimens are stored as observational evidence supporting phenotype interpretation. Measurements representing a molecular layer such as the genome, transcriptome, proteome, or karyotype are stored as Perspectives, including when those measurements arise from image-based assays. Spatial molecular assays therefore remain Perspectives, with coordinates stored as part of the molecular record rather than treated as purely phenotypic observations.

#### Event-linked perspective capture

The common Perspective representation is intentionally defined at the level of the molecular layer being profiled rather than the assay platform used to measure it. CLONEID does not require assay-specific ingestion routes because it does not ingest assays at the level of platform-specific file formats. Instead, it ingests a common event-linked Perspective representation: a biosample anchored to an Event, a set of inferred subpopulations with weights, and assay-specific feature matrices describing those subpopulations and optionally their members with spatial coordinates when relevant. This abstraction is broad enough to accommodate the shared core outputs of bulk clonal inference, single-cell clone assignment, and many spatial transcriptomic analyses after conversion into a common representation (Appendix 1.3), while method-specific auxiliary outputs such as trees and uncertainty may require additional metadata layers.

#### Event-linked phenotype capture and growth measurements

Phenotypic measurements were derived from event-linked observations recorded throughout the experiment. Bright-field images were uploaded and linked to their corresponding Events, and image-derived measurements, including segmentation-based features and cell counts were stored within the CLONEID record structure. Daily cell counts were used to generate fitted growth curves and growth-rate estimates across culture events. These longitudinal measurements were then queried directly from CLONEID for visualization and downstream comparison between adaptation histories.

### Derived phenotype quantities and processing assumptions

Several phenotype-associated quantities stored in CLONEID are derived rather than directly recorded (Table 2). For microscopy-based culture events, stored cell-count fields are estimated from segmented image fields and then scaled to the full vessel surface area using the recorded flask area. These values therefore represent whole-vessel approximations rather than direct counts of all cells in the culture vessel. Likewise, stored area-occupied values are extrapolated from segmented image fields to the full vessel area, and stored cell-size values are summary statistics derived from segmented cell areas rather than direct per-cell measurements. In the current implementation, cell size reflects a high quantile of per-image segmented-cell areas aggregated across images, rather than a direct mean size measurement.

Image-derived phenotype summaries may also be altered before final event-level values are stored. In the current pipeline, microscopy images are preprocessed before segmentation by default, including cell-line-aware gamma correction unless preprocessing is explicitly disabled. In addition, image-derived cell-count summaries may be filtered by an image-quality model and, when flagged, by manual exclusion of selected image positions before flask-level values are finalized. Microscopy segmentation parameters are also not fixed globally: cell-line-specific and, in some cases, lineage-specific parameter sets can be selected automatically, with a default parameter set used when no lineage-specific match is found.

### Event-history curation and modality-specific caveats

For workflows in which event identifiers encode ordered transitions, CLONEID uses two partially redundant sources of lineage information: the explicit parent–child linkage stored in the database and the implicit transition structure encoded in user-defined event identifiers. This redundancy is used as a validation mechanism rather than as an independent source of truth. In the current implementation, when an event identifier contains a single passage-designating token of the form `A<n>` (for example `A1`, `A7`, or `A24`), the insertion logic combines that token with event type and event date to infer the predecessor that would be expected under the transition sequence. More generally, the logic distinguishes <u>arrival-events</u>, which introduce a specimen into a new physical or contextual state, from <u>departure-events</u>, which precede or terminate such a transition. For arrival-events, the expected parent is a departure-event from the preceding stage; for departure-events, the expected parent is the corresponding arrival-event within the same stage that occurred earlier in time. Insertion is aborted only when this inferred parent is unique and conflicts with the supplied parent link. If the identifier does not support a unique inference, or if multiple candidate parents remain plausible, the system emits only a warning or message and does not automatically reject the record. This design aims to catch likely lineage-entry mistakes in long-term studies while avoiding overly rigid constraints on valid but ambiguously named records. Some event metadata may also be inherited by default: for example, departure events inherit flask identity from the parent event, and media information may be copied from the parent event unless specified explicitly.

Some non-microscopy modalities reuse phenotype storage fields with modality-specific meanings. In the current MRI path, a mask-derived volumetric quantity is stored through the same workflow that otherwise records microscopy-derived size summaries. These fields should therefore be interpreted in a modality-aware manner rather than assumed to represent the same physical quantity across all event types.

### Late karyotypic profiling as a molecular Perspective

At late stages of adaptation, samples from each of the six test trajectories were collected for karyotyping together with two parental controls. Karyotypic profiles were obtained for 28 *K*-selected cells, 61 *r*-selected cells and 32 control cells, for a total of 121 cells. Samples were collected after 27 serial culture events under *K*-selection and 55 serial culture events under *r*-selection. The cumulative area of segmented chromosomes was used as a surrogate for ploidy, and chromosome-level copy-number profiles were used for clustering and comparison of adaptation histories. In CLONEID, these karyotype-derived readouts were represented as late molecular Perspectives linked back to their event history and longitudinal phenotypic measurements.

### Preprocessing of genomic and karyotype data

#### scRNA-seq karyotype inference

For patient-derived analyses in the Appendix, we inferred changes in karyotype composition from single-cell transcriptomic data using Numbat, enabling comparison of inferred CNA states between primary and recurrent specimens.

#### Karyotype image processing

For karyotype images used in the density-selection demonstration and in Appendix analyses, karyotype panels were processed to segment individual chromosomes and assign chromosome labels. Karyotype images were required to contain chromosome groups arranged by standard karyotyping conventions with chromosome numbers annotated beneath each group. The analysis pipeline (i) detected annotated chromosome numbers and their image locations using optical character recognition, (ii) segmented chromosomes by thresholding and assigned each segmented chromosome to a chromosome group based on the detected label locations, and (iii) measured chromosome-level features including area (used as a ploidy surrogate in this work) and arm-level properties using morphological processing and skeletonization-derived endpoints.

### Mathematical modeling

For the Appendix patient cohort analysis, we used ALFA-K [19] as an external downstream modeling framework to infer local fitness landscapes for aneuploid karyotypes from single-cell karyotype data. CLONEID was used to store and organize the event-linked clinical and molecular records used as inputs to modeling (for example, imaging time points, biopsy Events and single-cell derived karyotype states), while fitness landscape inference and cross-validation were performed by ALFA-K.

### Interpretation of the demonstration analysis

Clustering of late karyotypic profiles separated cells according to their selection history. Most test-condition cells regained a copy of chromosome 21 relative to parental controls, and additional differences between *r*- and *K*-selected cells included loss of chromosomes 10 and 14 in the *K*-selected group. Karyotyping also identified tetraploidized cells enriched under *r*-selection, including whole-genome doubling in the r2 trajectory. Within CLONEID, these data were represented as late molecular Perspectives linked back to the corresponding event history and longitudinal phenotypic measurements.

## Acknowledgments

We thank Sue Grimes for her early contributions to the database design, which helped ensure its long-term scalability.

## Code availability

CLONEID is accessible through the web portal at dev.cloneid.org, which provides interactive access to the current CLONEID resource, including linked phenotypic, molecular, and specimen-context records. The portal also supports export of context-preserving subsets for downstream analysis. For users who wish to implement the CLONEID storage model within their own in-house database deployment, the corresponding cloneidR package is also available at https://github.com/thecloneredesignlab/cloneidR.git.

## Data availability

The data currently represented in CLONEID, including linked phenotypic, molecular, and specimen-context records, are accessible through the CLONEID portal at dev.cloneid.org. The portal supports interactive querying and export of context-linked subsets so that event history, specimen relationships, and associated molecular and phenotypic data remain connected during downstream reuse.

