## supplementary figures, tables and methods for "A longitudinal data framework for context-specific genotype-to-phenotype mapping"

#### APPENDIX

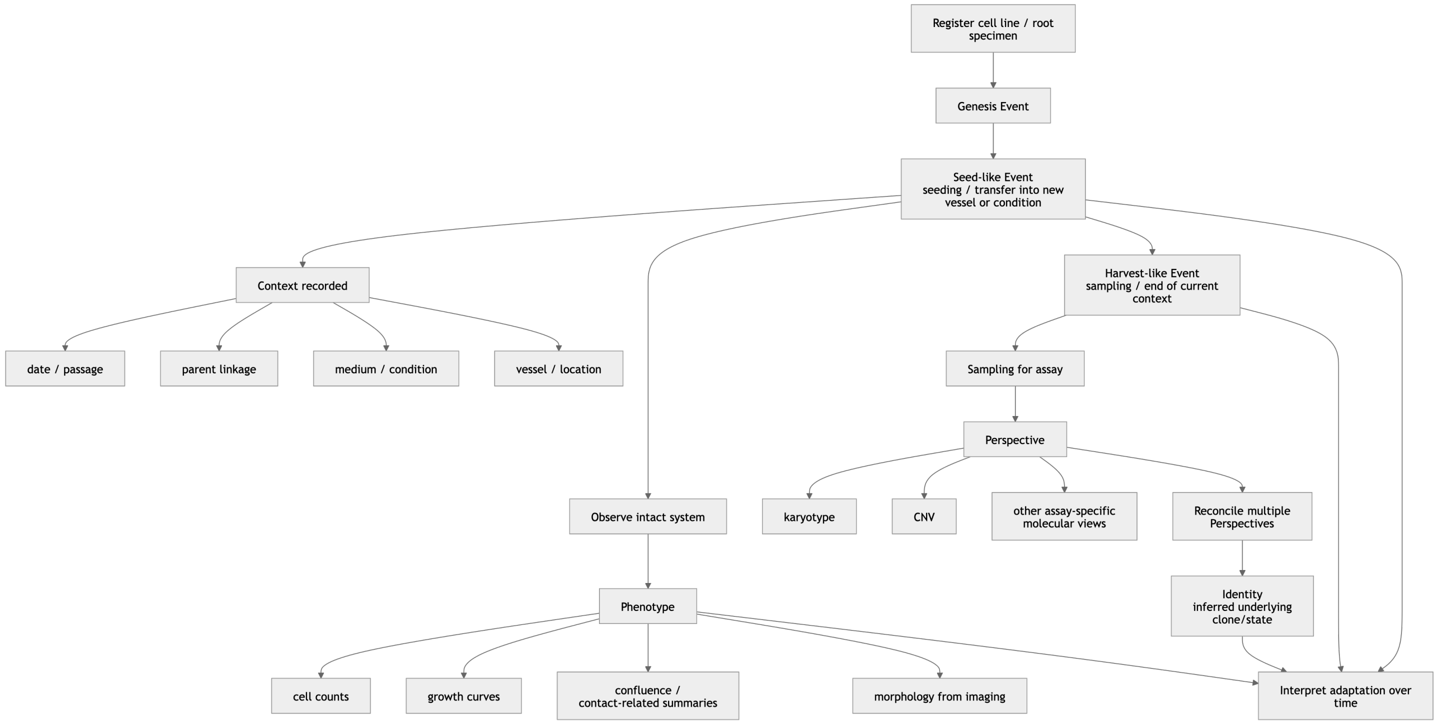


**Figure S1: Conceptual architecture of CLONEID.** CLONEID organizes longitudinal records around *Events*, which anchor specimen history. Each Event can support two complementary types of linked data. First, observation of the intact system yields *Phenotypes*, capturing behavior or state in context, such as growth, morphology, volume, motility, or treatment response. Second, sampling of material for assay yields *Perspectives*, which are assay-specific molecular views of the specimen, such as transcriptomic, copy-number, karyotypic, or spatial measurements. Because each Perspective is partial, multiple Perspectives can be reconciled into an *Identity*, representing an inferred underlying clone or state with preserved provenance. Together, Events, Phenotypes, Perspectives, and Identities support longitudinal interpretation of genotype–phenotype relationships across time and settings.

### 1 Interfaces and adoption

CLONEID is implemented as a defined SQL schema with a core functionality layer and complementary user-facing and programmatic interfaces. The web portal supports browsing, upload and subtree-based export of event-linked phenotypic and molecular records, whereas the public R package, (`https://github.com/theclonedesignlab/cloneidR`), supports a superset of portal functionality, including schema creation, event ingestion, Perspective ingestion, Identity reconciliation, querying, visualization and export. Across both interfaces, records are organized through the same event-based schema: users register Events, attach observational and molecular data as event-linked measurements and Perspectives, reconcile Perspectives into Identities and retrieve context-linked subsets for downstream analysis.

#### 1.1 User-facing interfaces and programmatic access

The CLONEID web portal provides two main user-facing modules together with a certified-user login. A *view* module allows users to search by cell line identifier or patient identifier and inspect specimen-event histories together with event-linked visualizations such as growth curves and time tables. An *upload* module supports ingestion of both event-linked phenotype records and multi-omics/genotype records through guided drag-and-drop workflows. In parallel, the public R package provides programmatic access to the same underlying schema for ingestion, reconciliation, querying, visualization and export. This dual interface design supports both interactive use by experimental researchers and scriptable analysis pipelines.

#### 1.2 Ingestion workflow and minimal required metadata

In practice, ingestion distinguishes registration of persistent entities, such as cell lines or patients, from registration of individual Events linked to those entities over time. Ingestion in CLONEID begins by registering a new Event with a persistent identifier and linking it to its source specimen or precursor Event when applicable. For relocation-like transitions, precursor linkage follows a specific arrival–departure rule rather than a generic "most recent prior Event" rule. An arrival Event is recorded after entry into a new physical location and links to the departure Event from the previous location that made that arrival possible. Conversely, a departure Event is recorded before exit from the current physical location and links to the arrival Event that placed the specimen in the location it is now leaving. This convention preserves the transition structure of specimen history across physical contexts. At a general level, each Event records an identifier, an event type, a timestamp, specimen relationships and controlled contextual fields whose interpretation depends on event type. In practice, this means that continuity-preserving Events can often be registered with relatively compact metadata, whereas transfer Events require explicit source–destination linkage and destination-specific context because the move itself may define a new evolutionary setting.

This scenario-neutral structure is used across clinical, in vivo and in vitro records (Fig. S2). For example, in vitro Events may include cell count, vessel context and media; in vivo Events may include injected cell count, strain and injection context; and clinical Events may include treatment context and imaging-derived summaries. Event-linked observations such as microscopy or radiology images are then attached to the corresponding Event and may be processed through versioned analytic modules, i.e. processing components that transform event-linked observations into derived intact-system descriptors while preserving provenance to the source data and originating Event. CLONEID defines this module role at the architectural level rather than requiring any specific analytic implementation, allowing current and future modules, including user-contributed ones, to extend support across imaging modalities and other event-linked observation types.

For portal-based ingestion of event-linked phenotype records, users select an event class and upload one or more associated image files. The current interface explicitly documents required or optional fields including a source event, timestamp, media/context field and, for seeding-style events, vessel type. To reduce manual relabeling errors, the Event identifier is derived from the uploaded file naming convention rather than being entered separately, helping ensure that phenotypic observations are attached to the correct Event record. Additional quantitative fields such as technician-estimated cell counts may also be entered when relevant. This workflow is designed so that derived phenotypic measurements remain anchored to the Event record.

##### 1.3 File conventions for multi-omics Perspectives

Multi-omics Perspectives are ingested using a defined folder convention that represents a biosample through a hierarchical clone-resolved molecular view. In the current implementation, users upload an unzipped folder containing a sample-level `.spstats` file together with one or more `.sps.cbs` files. The `.spstats` file lists inferred subpopulations for the biosample together with their cellular fractions. The sample-level `.sps.cbs` file stores an assay-specific feature matrix whose rows denote molecular features (first column `LOCUS`) and whose remaining columns denote inferred subpopulations. Optional additional `.sps.cbs` files provide lower-level member profiles for individual cells or members assigned to a given subpopulation. Perspective records may therefore represent molecular measurements at sample, subpopulation, or individual-cell resolution while remaining linked through a shared Event-anchored structure. In the current implementation, the uploaded folder is interpreted with respect to a Perspective type such as `GenomePerspective` or `TranscriptomePerspective`, and optional spatial coordinates may also be recorded when relevant.

A hard requirement of the current workflow is that the corresponding image/event record must already exist in CLONEID before a Perspective upload can be added to the database. In other words, molecular Perspectives are not ingested as free-standing records; they are attached to an existing event-linked specimen context. Following upload, the system reports stored profiles and visualizes clonal representation as an immediate ingestion check.

##### 1.4 Querying and context-bundled export

CLONEID supports interactive querying through the web portal by allowing users to search for a cell line identifier or patient identifier and then navigate the resulting event-history view. Node-level actions provide access to event-linked visualizations and available molecular views. For downstream reuse, the portal supports subtree-based export of a selected record subset. Export is performed from the selected subtree and includes the relevant entries together with their upstream Events and downstream Perspectives,

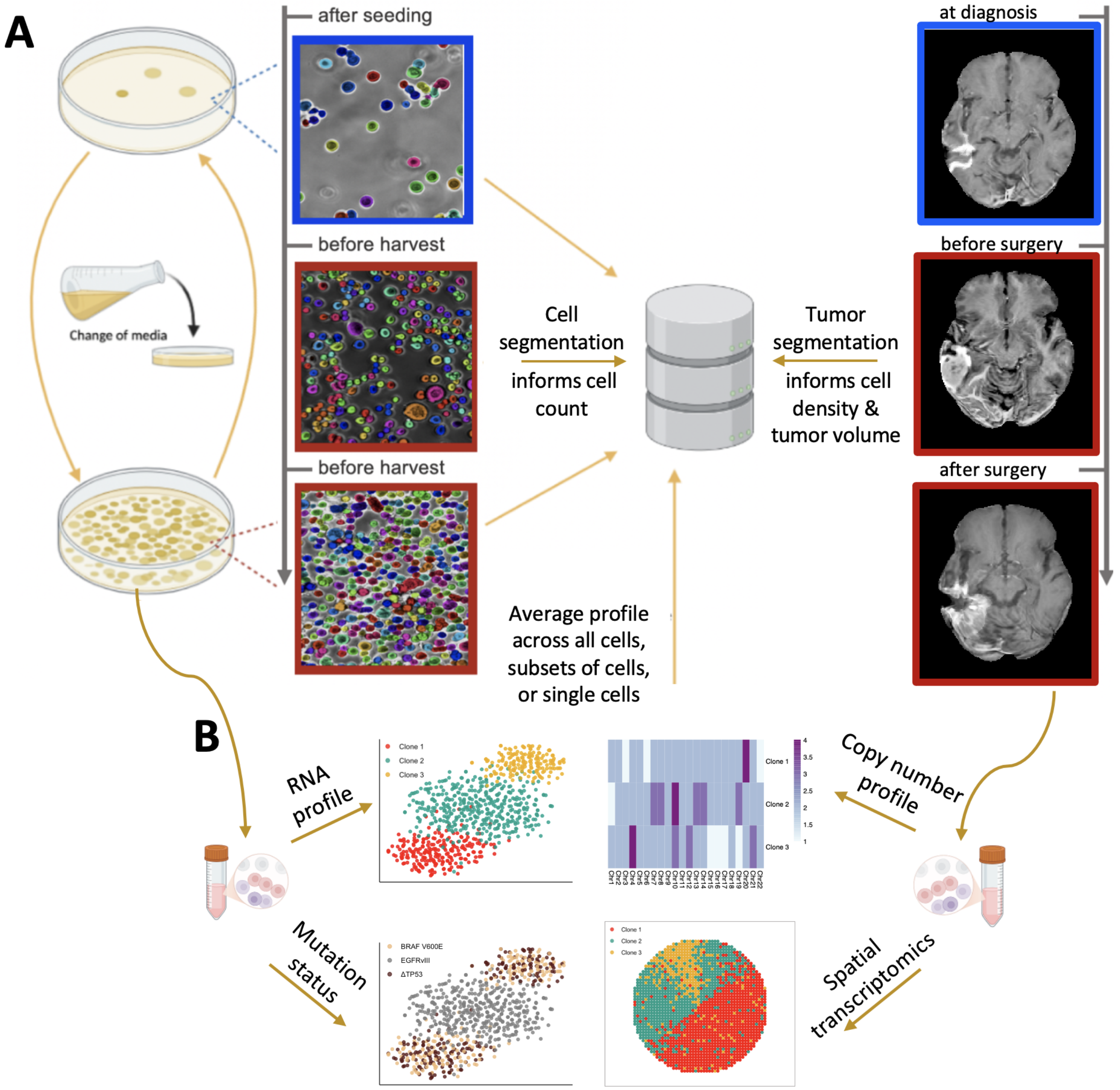


**Figure S2: CLONEID workflow and central components.** CLONEID is designed for multi-modal, longitudinal integration of phenotypic and genotypic data, explicitly linking genotype to phenotype across experimental or clinical timelines. (**A**) Phenotype module: each event (e.g., seedings, harvests, injections, resections) is recorded with precise timestamps linked to associated phenotype measurements such as cell counts, growth rates, or tumor volumes, together with event-context fields such as media conditions. Bright-field images (*in-vitro* NUGC-4, segmented with Cellpose) and clinical radiology scans from a glioblastoma patient illustrate the real-time nature of this phenotyping. (**B**) Genotype module: Single-cell and spatial assays identify clonal populations and map their molecular states back to events recorded in the phenotype module. From left to right: (i) t-SNE embedding of scRNA-seq colored by inferred clone; (ii) clone-specific copy-number heatmap across Chr 1–22; (iii) the same embedding colored by TISCC-seq informed [20] driver mutation (BRAF V600E, $\Delta TP53$, EGFRvIII); and (iv) spatial transcriptomic slice depicting patterns of clonal expansion. Panels A and B created with BioRender.com.

subject to a maximum traversal depth of 2 to limit excessively large downloads from high-level nodes.

The exported object is designed as a portable partial database. Subtree export produces a partial SQL database dump together with the associated imaging data. This bundled export preserves the contextual linkage between event history, phenotypic observations and molecular records, enabling downstream analysis in external environments without requiring direct access to the full CLONEID database.

**Table 2: Examples of stored or retrieved quantities in CLONEID that are derived rather than directly observed.**

| Stored field / output | Direct vs. derived | Current computation / rule | Key assumption / caveat |
|---|---|---|---|
| `cellCount`, `correctedCount` | Derived | Estimated from segmented image-field counts and scaled to the full vessel surface area using recorded flask area. | Represents an approximate whole-vessel count rather than a direct count of all cells present in the culture vessel. |
| `areaOccupied_um2` | Derived | Computed from segmented occupied area in image fields and extrapolated to vessel level. | Stored occupied area is a scaled whole-vessel estimate, not the raw segmented area from the imaged field alone. |
| `cellSize_um2` | Derived summary | Computed from segmented cell areas as an aggregated summary across images; in the current implementation it reflects a high quantile of per-image cell areas aggregated across images. | Does not represent a direct mean cell area measurement across all segmented cells. |
| Event-level microscopy phenotype summaries | Derived | Image fields may be excluded after quality-model assessment and operator review. | Final event-level values may depend on selective image exclusion. |
| Segmentation-derived phenotype values | Derived under lineage-aware parameterization | Segmentation parameters may be selected automatically from cell-line- or lineage-specific defaults, with fallback to a default parameter set. | Different branches of the same nominal cell line may be processed under different segmentation settings. |
| Event parentage | Inferred / validated | Parent-child links are validated against identifier-based passage structure and inferred probable ancestry. | Lineage relationships are constrained by naming conventions, not solely by free-form metadata entry. |
| MRI-derived phenotype field reuse | Derived, modality-specific | A mask-derived volumetric quantity may be stored through the same workflow otherwise used for microscopy-derived size summaries. | The meaning of a reused storage field can differ by modality and should not be assumed constant across event types. |

#### 1.5 Access modes and persistence

CLONEID distinguishes between exploratory upload workflows and persistent curation workflows. Through the web portal, uncertified users may test ingestion workflows and inspect derived outputs, whereas certified-user login is used for persistent data handling within the shared system. This access model provides a practical route for initial testing while preserving the integrity of curated database records.

#### 1.6 Storage model

CLONEID separates structured relational metadata from large binary and file-based artifacts. The relational database stores the records needed to reconstruct longitudinal context, including event-linked specimen history, assay-specific molecular views, inferred clone identities, and supporting provenance tables. In the manuscript, these records are described in terms of Event, Perspective, and Identity; in the current schema, some historical names remain, most notably `Passaging`, which implements event-linked specimen history. Molecular profiles are represented through `Perspective` and `Identity` records together with shared blob-backed payloads in `Loci`, so that measured views remain linked to their originating event records and sample-source metadata.

Large phenotype imaging and segmentation artifacts are stored outside the relational database. These include raw phenotype inputs together with derived outputs such as masks, annotations, detection results, and confluency measurements. In the current implementation, such artifacts are organized under durable S3-backed object-storage prefixes rather than embedded as SQL blobs, while the database retains the event and specimen relationships needed to interpret them. File names and object keys are anchored to event-linked record identifiers, allowing image-derived outputs to be re-associated with the correct specimen history during retrieval and export. This design keeps the relational schema focused on lineage, context, and provenance while allowing substantially larger binary assets to be stored and accessed independently.

This separation is operationalized through the phenotype storage layer, which reads and writes durable inputs and outputs through configured S3 roots while preserving linkage back to event-linked record identifiers. In practice, this provides a more scalable store for longitudinal imaging and derived segmentation outputs, and allows retrieval workflows to materialize only the files needed for a given task. To support context-preserving export, CLONEID can summarize the object-backed phenotype assets associated with a subtree of related events and combine those summaries with relational metadata to generate portable bundles and manifest templates for downstream selection and download.

This storage model supports provenance-preserving retrieval across longitudinal studies. Starting from an event-linked record, a user or downstream workflow can recover parent–child relationships from the relational database and then retrieve the associated assay views and phenotype artifacts without losing the surrounding history. As a result, phenotype measurements, molecular perspectives, and inferred clone identities can be stored in different physical layers while still being reconstructed together as a coherent longitudinal record. This separation of storage layers is what allows CLONEID to support both interactive portal retrieval and reproducible downstream analysis.

#### 2 Database content and current resource coverage

As of this publication, the CLONEID database contains **6,378 event-anchored specimen records** organized into 17 retained case histories spanning clinical, in vivo and in vitro datasets, with data types ranging from bulk and single-cell DNA/RNA-derived molecular perspectives to image-derived phenotypes. More than 30,000 microscopy, digital pathology and medical images are stored with their associated event context and specimen relationships (Fig. S3A).

The multi-omics component stores measurements from 103,973 individual cells and 357 distinct clone-level objects, organized into modular layers (for example, *GenomePerspective* and *TranscriptomePerspective*) that can be linked to clonal identities through Perspective and Identity records (Fig. S3B). In addition to interactive querying, CLONEID supports exporting context-linked subsets so that events, phenotypes and molecular views remain linked for downstream reuse.

#### 3 Additional demonstration analyses

##### Use Case 2: Predicting cellular plasticity from a snapshot of the transcriptome

Four gastric cell lines in the CLONEID database (HGC-27, NCI-N87, SNU-16 and SNU-668) were propagated for 40 or more serial culture events, representing months of continuous culture. Event-linked images were analyzed to quantify cell numbers over time, and growth models were fit to these longitudinal trajectories to summarize changes in growth rate across the course of culture adaptation (Fig. S4A,B).

To ask whether genetic structure could predict divergence in growth trajectories, copy-number-based phylogenetic trees were constructed from single-cell transcriptomic snapshots using CONICs [21]. For each tree, pairwise coalescence times were computed for sequenced cells and compared between adapting and non-adapting lines (Fig. S4C,D). This analysis illustrates how event-linked phenotypes stored in CLONEID can be paired with later molecular readouts to support retrospective, hypothesis-generating analyses.

##### Use Case 3: Inferring karyotype fitness landscapes from patient data

We used CLONEID to organize multimodal clinical and molecular records from a cohort of 11 glioblastoma (GBM) patients with matched primary and recurrent tumor specimens [22, 23]. The stored records included event-linked imaging time points, biopsy Events, and single-cell derived karyotype states (Fig. S5A,B). We then applied ALFA-K to infer patient-specific karyotype fitness landscapes from these single-cell karyotype dynamics [19]. For example, in patient P30, ALFA-K inferred fitness values across 28,405 unique karyotypes (Fig. S5C) based on the dynamics of karyotypes observed in the input data

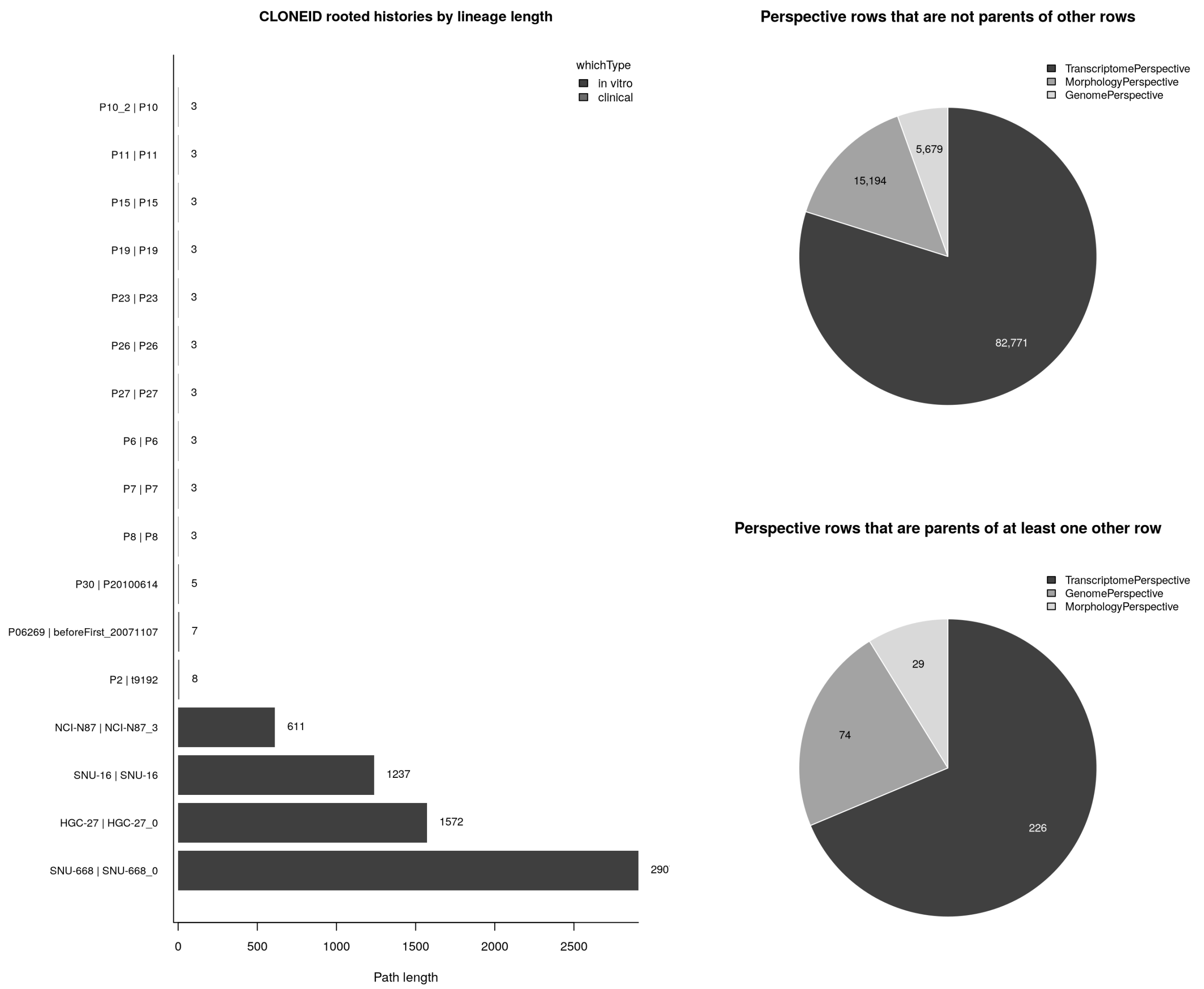


**Figure S3: Current CLONEID resource coverage and representation across histories and molecular layers.** Left, summary of rooted histories currently represented in CLONEID, showing lineage length for the 24 longest retained histories, with all remaining shorter histories collapsed into "others." Right, composition of Perspective records by 'whichPerspective', shown separately for single-cell entries that are not the parent of any other Perspective record (top) and clone-level entries that serve as the parent of at least one other Perspective record (bottom).

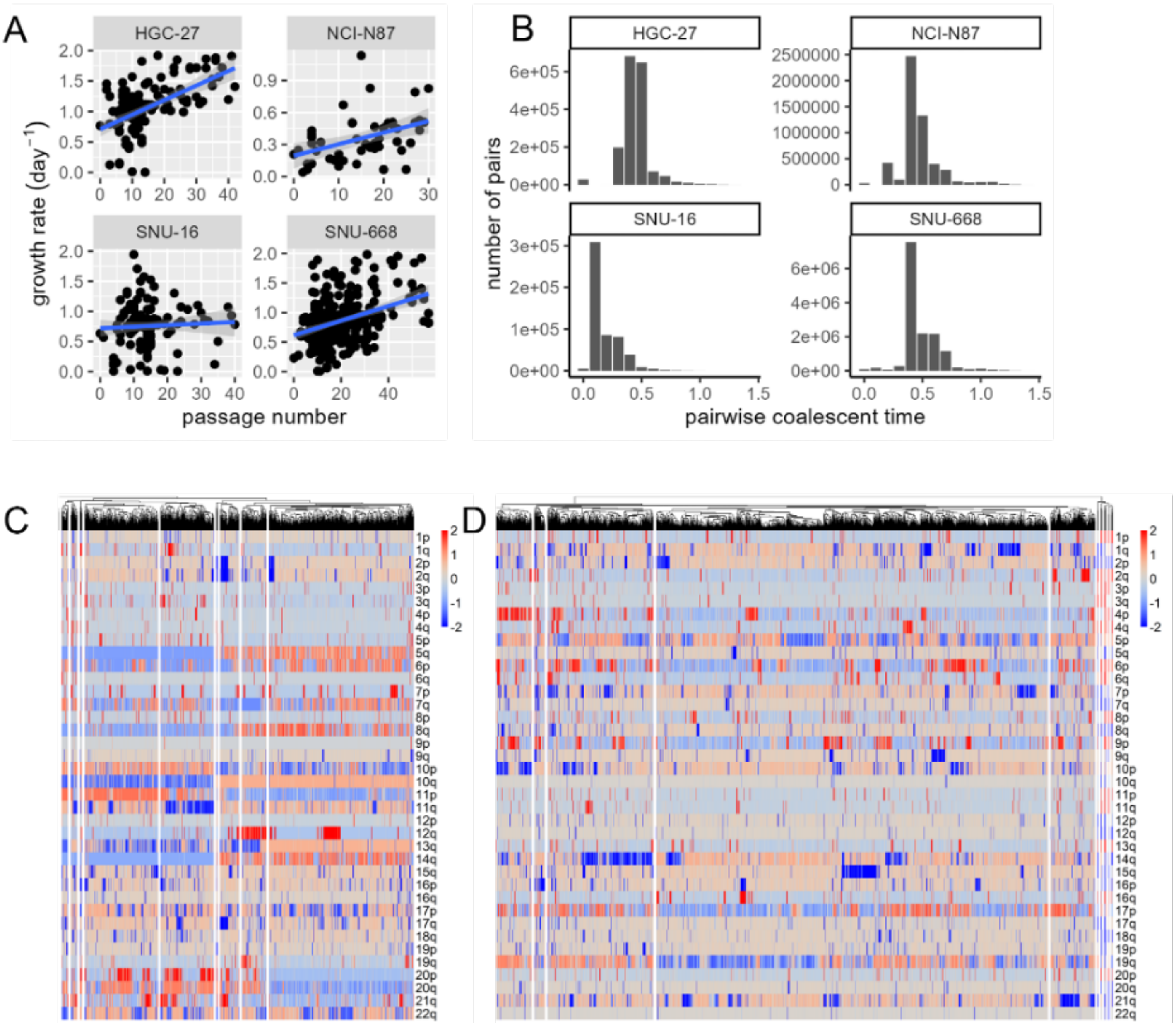


**Figure S4: Predicting cellular plasticity from a snapshot of the transcriptome**. (A) Event history data were used to reconstruct specimen graphs, and logistic growth models were fit to cell-count trajectories from culture events with at least three time points to estimate intrinsic growth rates. These growth rates were plotted against event number to reveal temporal changes in proliferative dynamics. (C,D) CONICSmat was applied to scRNA-seq data to infer large-scale copy-number variations from normalized expression matrices. After scaling and thresholding the data, hierarchical clustering produced a copy-number-based phylogenetic tree, from which pairwise coalescence times were computed. Trees and coalescence-time structure were compared across adapting and non-adapting lines.

(Fig. S5B). Cross-validation showed strong concordance for 9 out of 11 cases, which were included in downstream analyses.

Across these landscapes, we observed both shared and patient-specific selection pressures: among 203,064 unique inferred karyotypes, a minority occurred in more than one patient landscape, and the correspondence of inferred fitness across patients varied, consistent with context-dependent CNA fitness effects.

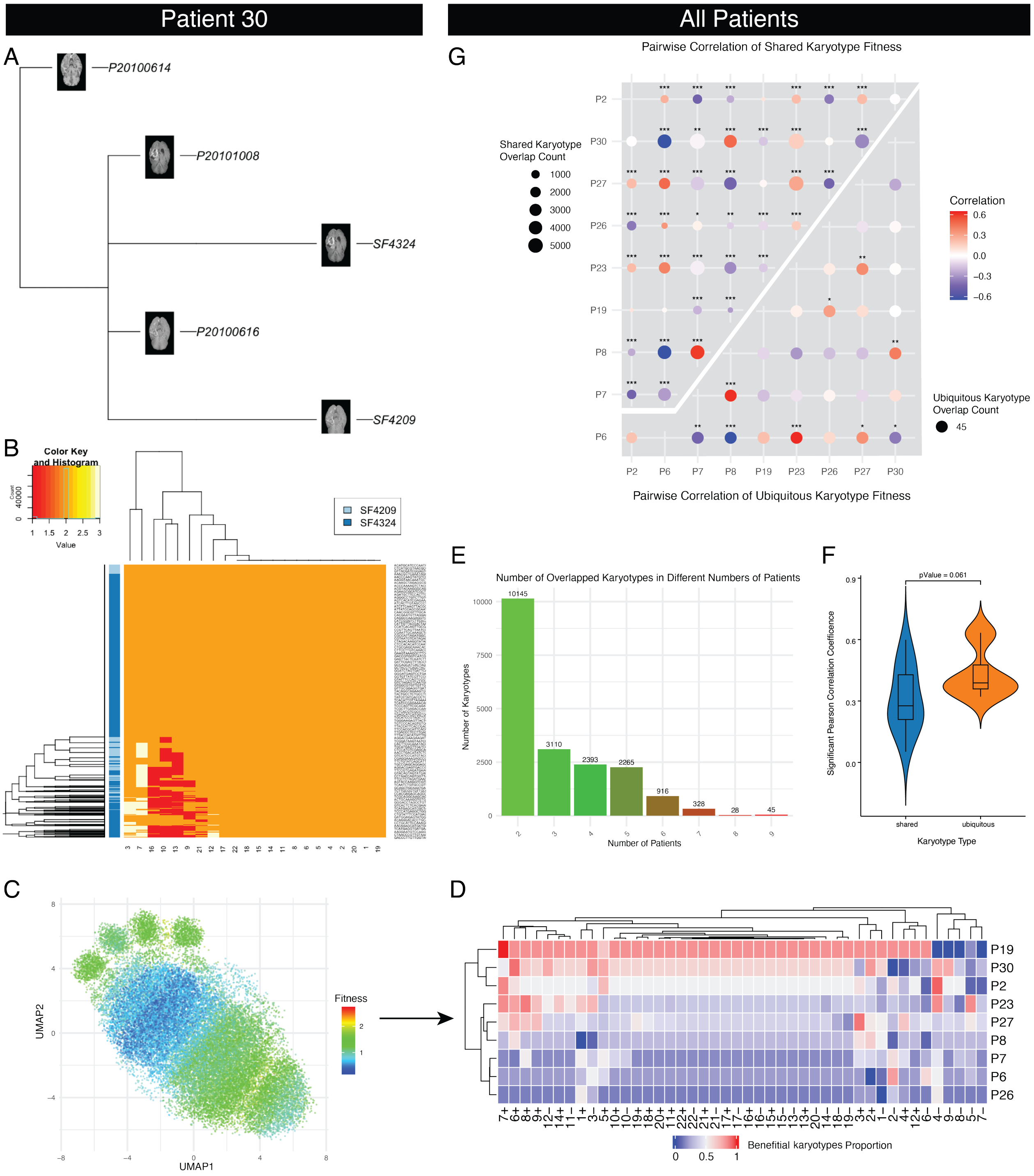


**Figure S5:** Multimodal analysis of a Glioblastoma patient through CLONEID informs a mathematical model of karyotype evolution. (A) MRI data from three time points were viewed through CLONEID's phenotype module; biopsies were taken during surgery at two time points (prefix SF*). (B) Karyotype composition derived from scRNA-seq of the two biopsies. 4,471 cells (rows) were sequenced in total and were used as input to a mathematical model for inferring karyotype fitness landscapes. (C) UMAP representation of 28,405 unique karyotypes that make up the charted fitness landscape for P30. Each karyotype is color-coded by its inferred fitness. (D) For each CNA, the proportion of karyotypes in which that CNA increases fitness is shown. (E) Summary of karyotype overlap across inferred fitness landscapes. (F) Pearson correlations between fitness values of ubiquitous karyotypes across patient pairs. (G) Pairwise correlations of fitness values for shared karyotypes (bubble chart above diagonal) and ubiquitous karyotypes (bubble chart below diagonal).
